# Early ctDNA kinetics as a dynamic biomarker of cancer treatment response

**DOI:** 10.1101/2024.07.01.601508

**Authors:** Aaron Li, Emil Lou, Kevin Leder, Jasmine Foo

## Abstract

Circulating tumor DNA assays are promising tools for the prediction of cancer treatment response. Here, we build a framework for the design of ctDNA biomarkers of therapy response that incorporate variations in ctDNA dynamics driven by specific treatment mechanisms. We develop mathematical models of ctDNA kinetics driven by tumor response to several therapy classes, and utilize them to simulate randomized virtual patient cohorts to test candidate biomarkers. Using this approach, we propose specific biomarkers, based on ctDNA longitudinal features, for targeted therapy, chemotherapy and radiation therapy. We evaluate and demonstrate the efficacy of these biomarkers in predicting treatment response within a randomized virtual patient cohort dataset. These biomarkers are based on novel proposals for ctDNA sampling protocols, consisting of frequent sampling within a compact time window surrounding therapy initiation – which we hypothesize to hold valuable prognostic information on longer-term treatment response. This study highlights a need for tailoring ctDNA sampling protocols and interpretation methodology to specific biological mechanisms of therapy response, and it provides a novel modeling and simulation framework for doing so. In addition, it highlights the potential of ctDNA assays for making early, rapid predictions of treatment response within the first days or weeks of treatment, and generates hypotheses for further clinical testing.

## 1 Introduction

Circulating tumor DNA (ctDNA) assays – which detect cell-free DNA fragments released by tumor cells into the bloodstream – are a highly sensitive, noninvasive approach to assessing tumor burden and genomic profiles without requiring biopsy or surgical resection. Indeed, ctDNA analyses have been proposed for a variety of clinical purposes, including early cancer detection, detection of tumor recurrence, general surveillance of treatment response, and assessment of minimal residual disease (e.g. [28, 11, 35, 30, 3, 23, 32, 16]). Serial ctDNA kinetics have also been explored as a possible predictor of response to treatment in a variety of cancer types, such as colorectal cancer [36, 10, 33, 8, 25] and non-small cell lung cancer [1, 38, 2, 37]. Some features of ctDNA longitudinal dynamics, such as the existence of transient peaks in ctDNA levels, rapid clearance rates during therapy, or lower ctDNA levels at baseline or a month after therapy, have been correlated with treatment efficacy [30, 24, 27, 9, 31, 33, 14]. Specific features of these early data vary greatly between different treatment types [30], For instance, for ctDNA primarily shed during cellular apoptosis, treatments that induce higher levels of inflammation, turnover and apoptosis may produce larger initial spikes in ctDNA, as compared to primarily cytostatic therapies [22]. suggesting that any longitudinal biomarkers designed for interpreting early ctDNA dynamics should be specific to therapeutic class and mechanism of action.

Currently, standardized clinical approaches to the collection of ctDNA data or their interpretation for therapeutic decision-making are lacking. A majority of existing studies utilizing ctDNA as clinical biomarkers have focused on examining ctDNA levels or clearance at a single time point. When investigated serially, the frequency of ctDNA collection can vary widely and is largely clinician-dependent, and the collection of baseline samples prior to treatment initiation is similarly inconsistent. In some cases, analyses are performed on samples acquired during and/or post treatment, usually at time points several months apart, including at time of disease progression. Since the production and clearance kinetics of ctDNA in the bloodstream occur on relatively faster time scales (hours, minutes) [4, 35], this spacing may fail to capture valuable information about treatment efficacy available from the early stages of therapy induction. Sanz-Garcia et al. [30] provide an excellent review of the existing data on ctDNA kinetics and biomarker trials under various types of treatment, and Table 1 shows a partial summary of existing studies on ctDNA biomarkers. These studies demonstrate significant potential in the use of ctDNA analyses for prognostic predictions, but also highlight the need for better standardization of collection techniques, increased prediction accuracy and an improved understanding of how early ctDNA dynamics vary across treatment types.

**Table 1:** A partial summary of ctDNA biomarker analysis in targeted therapy, chemotherapy, and radiotherapy.

| Author(s) | Treatment type | $n$ | Time points | Biomarker definition | Efficacy |
| --- | --- | --- | --- | --- | --- |
| Parikh et al. [24] | Targeted therapy | 138 | Baseline, 2, 4, 8, and 16 weeks | Decrease from baseline to wk 4. | Predicted clinical benefit at 90% specificity and 60% sensitivity |
| Vidal et al. [34] | Chemotherapy and targeted therapy | 100 | Baseline and C3 (week 4-6) | Decrease from baseline | Correlation with progression free survival (hazard ratio 0.23, $p = 0.001$) |
| Magbanua et al. [16] | Chemotherapy | 295 | Baseline, 3 and 12 weeks, post-treatment | Concentration, clearance, and others | Early clearance correlated with pathologic complete response (odds ratio 13.06, $p = 0.0002$) |
| Garlan et al. [8] | Chemotherapy | 82 | Baseline, 2 and 4 weeks | Concentration at baseline, decrease from baseline | Dividing patients into “good” and “bad” ctDNA responders. Good responders had better objective response rate (41.3%) than bad responders (0%) |
| Tie et al. [33] | Chemotherapy | 53 | Baseline, day 3, and day 14-21 | $\geq 10$-fold reduction from baseline | 74% with $\geq 10$-fold had radiologic response, compared to only 35% of patients with lesser reductions |
| Leung et al. [13] | Radiotherapy and chemo-radiotherapy | 107 | Baseline, 4 weeks, and post-treatment | Detectable ctDNA at 4 weeks | Correlated with worse progression free survival (hazard ratio 4.05, $p < 0.0001$) |
| Chera et al. [5] | Chemo-radiotherapy | 103 | Baseline, 1, 2,3, and 4 weeks | Favorable clearance profile: high baseline and $>95\%$ clearance by day 28 | 19 out of 67 patients had favorable clearance profiles: none had persistent or recurrent disease. 35% of patients with unfavorable profiles had persistent or recurrent disease ($p = 0.0049$). |
| Lv et al. [15] | Radical induction chemotherapy and chemo-radiotherapy | 673 | Once per cycle for four cycles | Used supervised statistical clustering to create four prognostic phenotypes | Clusters associated with different risks of tumor relapse |

In this study, we explore the potential of early ctDNA dynamics as a prognostic indicator of treatment efficacy, using mathematical modeling as a conceptual tool. Specifically, we develop mathematical models for ctDNA dynamics under targeted therapy, chemotherapy and radiotherapy, and leverage these models to explore how ctDNA kinetics vary between therapy classes. These results build upon a growing literature on mathematical models of ctDNA dynamics. For example, Avanzini et al. [3] introduced a mathematical model of ctDNA dynamics in untreated tumors and estimated the increase in detection lead time afforded by ctDNA analyses over imaging. In another work, Khan et al. [10] explored the potential of ctDNA to detect RAS mutants as an early signal of resistance in colorectal cancer. These mathematical models, as well as other computational studies [26, 2, 6] have demonstrated the tremendous potential of using observed ctDNA dynamics to extract insights about tumor biologic processes and response to treatment. Here, using simulated virtual patient cohorts, we develop dynamic biomarkers predictive of treatment response using early, frequent ctDNA sampling. We demonstrate that these biomarkers, based on mechanistic biological principles, perform equally as well as standard neural network approaches while retaining mechanistic interpretation. These modeling results suggest that frequent ctDNA collection in the initial stages of treatment may provide a critical early indicator of whether treatment will ultimately be successful.

### 2 Impact of therapeutic mechanisms on ctDNA dynamics

We first explore how various therapeutic mechanisms of action may drive differences in ctDNA dynamical features. In particular, we develop mathematical models of ctDNA kinetics arising from tumor cell population responses to targeted therapy, chemotherapy and radiotherapy, and examine how these mechanisms drive distinct features in ctDNA data.

### 2.1 Targeted therapy model

We consider a mechanistically-motivated mathematical model of ctDNA shedding under cytotoxic targeted therapy, as described in Figure 1b. Therapies often fail due to the emergence of drug-resistant cell subpopulations within a tumor; thus, we incorporate both drug-sensitive and drug-resistant subpopulations within the model. Each population evolves as a stochastic birth-death process, in which cells divide and die stochastically with exponentially distributed waiting times governed by their respective birth and death rates, which may vary according to cell type and drug concentration. Specifically, the drug-sensitive population has birth rate *b*_*s*,1_, death rate *d*_*s*,1_ and net growth rate *λ*_*s*,1_ *≡ b*_*s*,1_ *− d*_*s*,1_ in the absence of drug; it has birth rate *b*_*s*,2_, death rate *d*_*s*,2_ and net growth rate *λ*_*s*,1_ *≡ b*_*s*,2_ − *d*_*s*,2_ in the presence of drug. We assume that this population on average expands in the absence of therapy and declines in the presence of drug (i.e. *λ*_*s*,1_ > 0 *> λ*_*s*,2_). The drug-resistant population has birth rate *b*_*r*,1_, death rate *d*_*r*,1_ and net growth rate *λ*_*r*,1_ *≡ b*_*r*,1_ − *d*_*r*,1_ in the absence of drug; it has birth rate *b*_*r*,2_, death rate *d*_*r*,2_ and net growth rate *λ*_*r*,1_ *≡ b*_*r*,2_ *− d*_*r*,2_ in the presence of drug. We assume that therapy can impact the growth of drug-resistant cells, but that the drug-resistant population increases on average, both in the presence and absence of drug (i.e. *λ*_*r*,1_ ≥ *λ*_*r*,2_ > 0).

**Figure 1:**
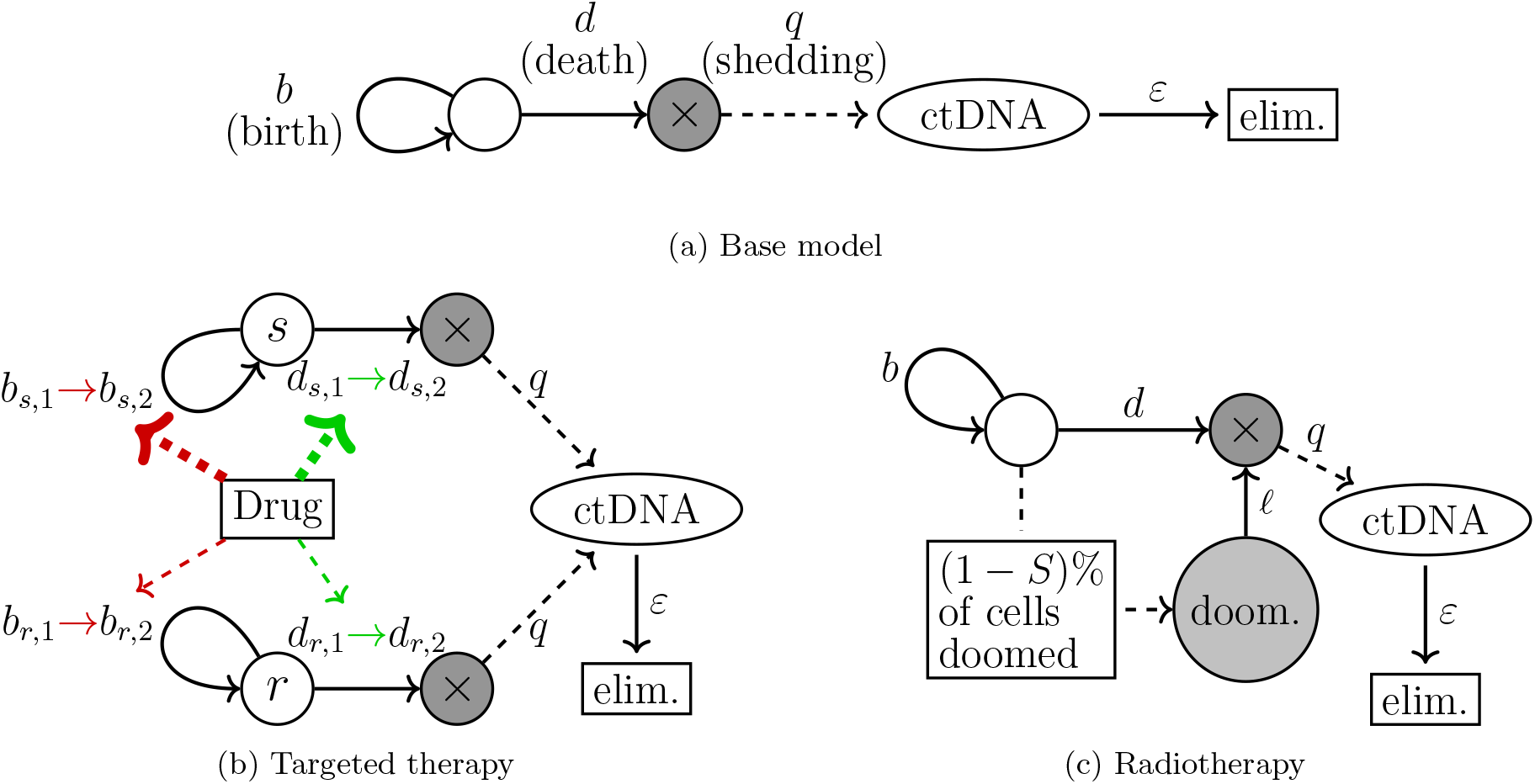
Schematics of models for ctDNA kinetics: baseline without treatment, under targeted therapy, and under radiotherapy. **(a)** Base model: Without treatment, the tumor population is modeled as a stochastic birth-death process, starting with an initial population with birth rate *b* and death rate *d*. For each cell death, one hGE of ctDNA is released with probability *q*. ctDNA is eliminated from the bloodstream at rate *ε*. This model was introduced in [3]. **(b)** Targeted therapy model: Initial tumor population consists of a mixture of sensitive and resistant cells, modeled as stochastic birth-death processes with (untreated) birth and death rates *b*_*s*,1_, *d*_*s*,1_ and *b*_*r*,1_, *d*_*r*,1_, respectively. Under therapy, the birth and death rates of the sensitive and resistant populations are *b*_*s*,2_, *d*_*s*,2_ and *b*_*r*,2_, *d*_*r*,2_, respectively. **(c)** Radiotherapy model: Initial population of cells with birth rate *b*, death rate *d*. For each dose of radiotherapy, *S* is the proportion of cells that survive each treatment. The remaining (1 *− S*) proportion of cells are lethally irradiated cells which can no longer divide but experience mitotic catastrophe and die at rate *𝓁*, at which point they release ctDNA with probability *q*.

Similarly to [3], to model the process of ctDNA production we assume that during each cell death from either population, one human genome equivalent (hGE) of ctDNA is shed into the bloodstream with probability *q*. The ctDNA is eliminated from the bloodstream at random exponential rate *ε*. We assume in this model that the modeled ctDNA tracks a generic tumor genomic marker that does not distinguish whether the ctDNA originated from the drug-sensitive or drug-resistant subpopulation; future work will consider more detailed models of individual mutational frequencies. In our treatment model, cytotoxic drug is applied continuously starting at time 0 and may alter the birth and death rates of the tumor subpopulations. Under this model, we derive summary statistics for the abundance of ctDNA under treatment which are available in Appendix 6.3.

Figure 2a shows an example simulation of a tumor population that initially consists primarily of drug-sensitive cells, but also harbors a small resistant subpopulation. Upon application of treatment at time 0, the tumor population decreases and is accompanied by a sharp increase in ctDNA followed by a gradual decrease in ctDNA below baseline levels; this transient peak arises due to the sharp increase in cell apoptosis at the start of therapy application, resulting in a significant amount of released ctDNA. If the initial resistant subpopulation frequency is increased, the size of the initial peak is decreased 2c. Similarly, increasing drug effectiveness on the sensitive cell population results in higher initial peaks (Figure 2d), suggesting that the presence of transient peaks followed by a gradual decrease, as well as transient peak height, may be predictive of responsiveness in targeted therapy. Although datasets with sufficiently frequent, early samples of ctDNA are too rare to fully parametrize and validate this model, we fitted our model to evaluate consistency with one existing longitudinal dataset. Riediger et al. [27] examined daily ctDNA levels in a patient with non-small cell lung cancer treated with tyrosine kinase inhibitors (TKIs). The patient responded well to treatment and exhibited an 11-fold peak in ctDNA levels at 26 hours followed by a subsequent decrease. An example fitting of our derived expectation for ctDNA under the targeted model with this data is shown in Figure 2b. The same pattern was observed by Husain et al. [9], who obtained daily urine ctDNA samples from NSCLC patients treated with osimertinib, a second line anti-EGFR TKI all of whom exhibited clinical benefit. Eight patients had detectable ctDNA with T790M mutations, and all of them exhibited a spike in the first week of therapy followed by a large and sustained decrease in the following weeks.

**Figure 2:**
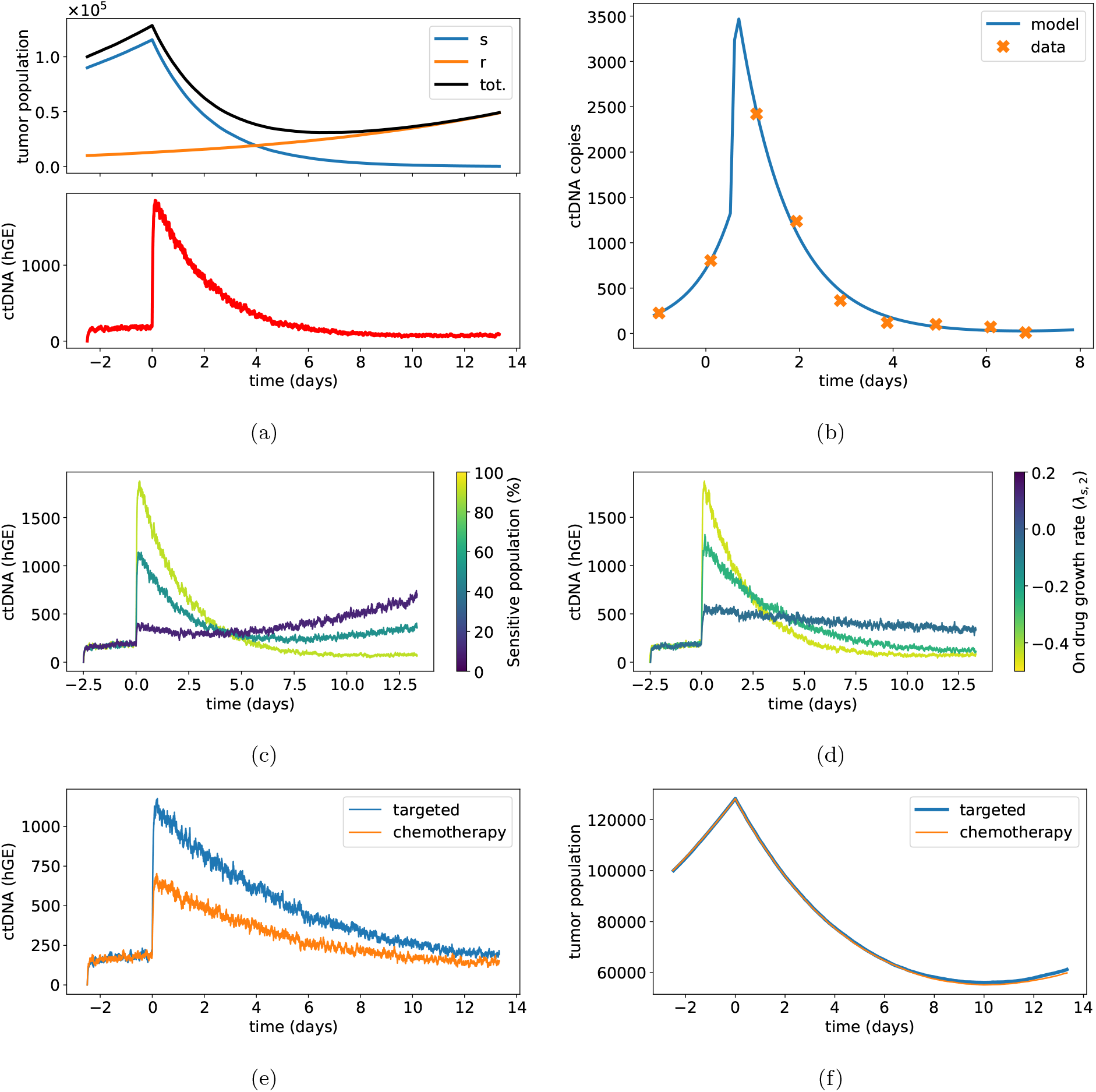
Exploration of targeted therapy model. **(a):** Example simulation for the targeted therapy model, initialized with 10^5^ total cells, 90% of which are drug-sensitive. The ctDNA shedding rate is set to *q* = 1 and the elimination rate is set to *ε* = 33 per day. Off drug, the birth rate is *b*_−,1_ = 0.15 per day and death rate is *d*_−,1_ = −0.05 per day for both drug-sensitive and drug-resistant cells. Treatment is started when the tumor increases in population by 25%. Upon treatment, the birth and death rates for the drug-sensitive cells become *b*_*s*,2_ = 0.1, *d*_*s*,2_ = 0.55 while the rates for the drug-resistant cells are unchanged. **(b):** Example linear regression fitting of our targeted therapy model to ctDNA data from a NSCLC patient treated with afatinib obtained by [27]. The blue line is the expected ctDNA levels 𝔼[*C*(*t*)] (see Appendix 6.3) produced by the best fit parameters and the orange crosses are ctDNA samples from patient data. **(c), (d):** Example ctDNA simulations for varied initial proportions of sensitive cells and drug efficacy. (c) shows simulations for proportions of 90%, 50%, and 10% sensitive cells. (d) shows simulations for varied death rate on drug resulting in values for *λ*_*s*,2_ of −0.45, −0.25, and 0.05 per day. The initial parameters of the simulations are the same as in (a) otherwise. **(e), (f):** A comparison of ctDNA levels from simulations of the targeted therapy model with *d*_*s*,2_ = 0.32 and the chemotherapy model with *K*_*s*_ = 0.9. Both simulations have the same net growth rates on and off drug *λ*_*s*,1_ = 0.1 and *λ*_*s*,2_ = −0.17.

#### 2.2.1 Adaptation for modeling chemotherapy

Note that the targeted therapy model can also be adapted to model ctDNA kinetics under chemotherapy. Under the chemotherapy setting of this model, a birth event of a sensitive cell has probability *K*_*s*_ of becoming a death event instead and the birth event of a resistant cell has probability *K*_*r*_ of converting to a death.This results in birth and death rates

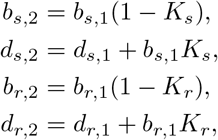

for the sensitive and resistant cells. Thus, the chemotherapy model is a special case of the targeted therapy model where the changes in the birth and death rates are dictated by *K*_*s*_ and *K*_*r*_.

Figure 2e shows a comparison of the difference in ctDNA levels between cytotoxic targeted therapy and chemotherapy. In both simulations, parameters were chosen to have the same net growth rate on and off drug so that the therapies are equally effective. However, we note that although the tumor population dynamics are effectively identical (Figure 2f), the ctDNA kinetics are markedly different, with the chemotherapy simulation exhibiting a significantly smaller peak than the targeted therapy. Intuitively, the explanation for this difference is that the chemotherapy model converts birth events to deaths which exerts control on the net growth rate by both decreasing the birth rate and increasing the death rate. On the other hand, cytotoxic targeted therapy primarily increases the death rate alone, leading to a greater ctDNA peak.

The behavior of the chemotherapy model is otherwise very similar to that of the targeted therapy model, with responsive patients exhibiting ctDNA peaks followed by gradual declines. These patterns are also observed clinically. Tie et al. [33] examined ctDNA in metastatic colorectal cancer (mCRC) patients receiving standard first-line chemotherapy. Four patients exhibited a moderate spike at day 3 (≤3-fold), followed by a rapid decline. Three of these patients experienced a tumor reduction of over 20%, leading the authors to hypothesize that the spike could reflect increased cell death from successful treatment.

### 2.2 Radiotherapy

We next developed a model of ctDNA dynamics driven by tumor responses to radiation therapy delivered in fractionated doses. The tumor population *N* (*t*) is modeled by a stochastic birth-death process as in our base model, with each death accompanied by probability *q* of shedding 1 hGE of ctDNA. Fractionated radiation treatments are applied at times {*t*_1_, … , *t*_*n*_}. The effect of each dose is modeled with a linear quadratic cell kill model where the survival probability of a cell following treatment with dosage *D* Gray is given by 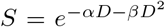[7]. This well-accepted model has been empirically validated in many studies, see, e.g. review [12]. For each dose, *SN* (*t*_*i*_) of the cells survive and are unaffected and (1 *− S*)*N* (*t*_*i*_) become lethally rradiated cells. These lethally irradiated cells can no longer divide and die off at rate *𝓁*. Rostami et al. [29] observed a delayed release of ctDNA between 96h and 144h after radiation treatment. Thus we set *𝓁* to be approximately 0.15 − 0.30 per day. Derivations of summary statistics for the abundance of ctDNA under radiotherapy treatment are available in Appendix 6.4.

Figure 3a shows an example simulation of the radiotherapy model with four cycles of weekday treatments starting on day 7. We observe that under this model, the ctDNA dynamics exhibit a different shape from the targeted therapy model; here, levels gradually peak several days after the first treatment date and exhibit a slow decline afterwards that tracks the tumor burden decline. Dynamical features arising from the fractionation schedule (vertical grey lines) are observable in finely sampled data. Figure 3c shows ctDNA kinetics for simulations with varied values of *S*, the survival fraction. Increased treatment efficacy results in a larger increase over the first week of treatment and a greater decrease in the second week of treatment. The parameter *𝓁* is associated with lag time between irradiation and cell death / ctDNA release. To investigate the model dynamics for varying lag times, Figure 3d shows simulations for varied mean lag times *𝓁*^−1^. While behavior during the second week of treatment and beyond does not change significantly, lower lag times result in increased ctDNA during the first week of treatment.

**Figure 3:**
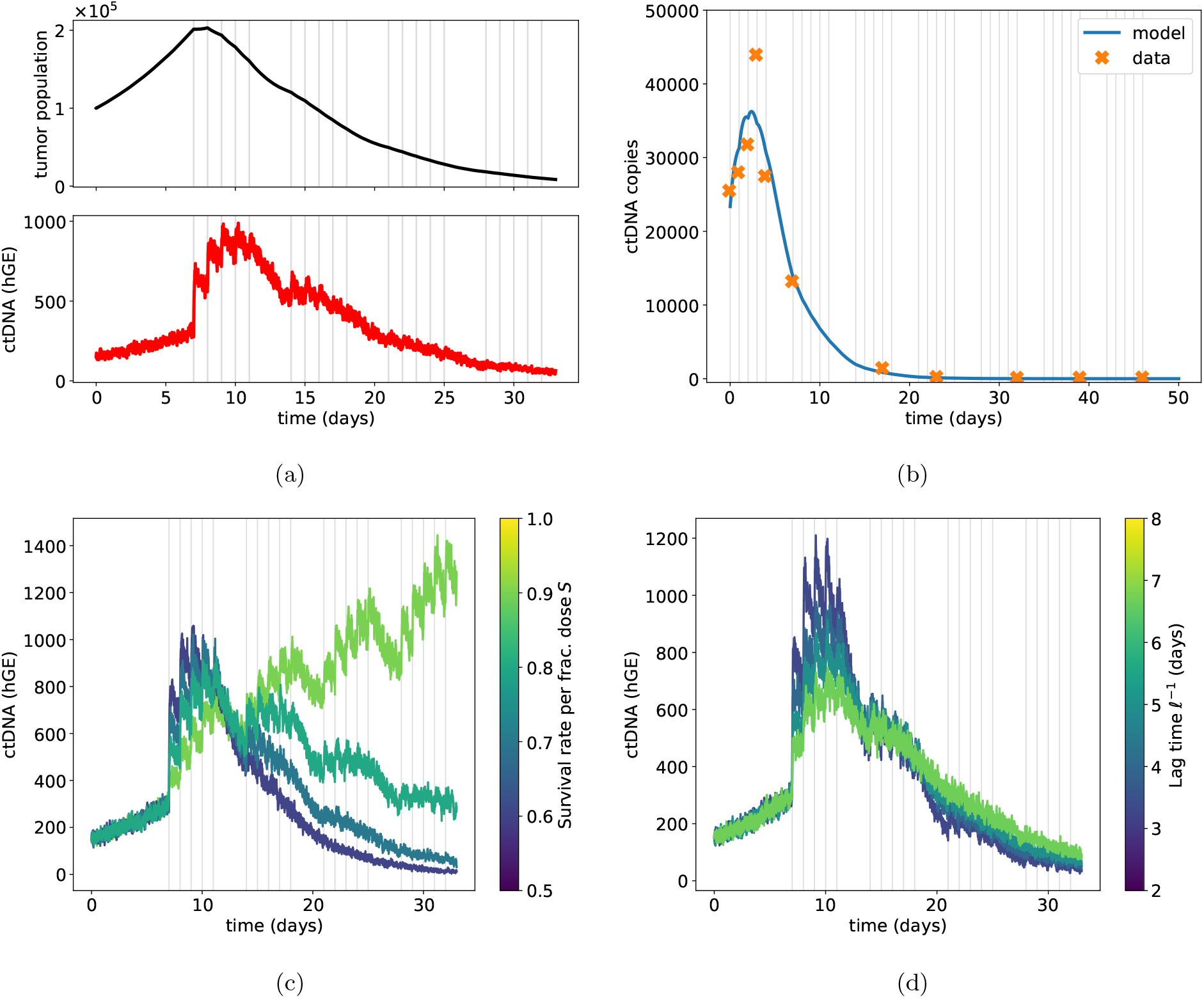
Exploration of radiotherapy model. **(a):** Example simulation starting with 10^5^ initial cells, birth and death rates *b* = 1.5, *d* = 0.05, elimination rate *ε* = 33, and shedding rate *q* = 1. Fractionated radiotherapy doses are applied in a 5 days on, 2 days off schedule starting on day 7. Each dose has survival rate *S* = 0.7. The lag parameter is *𝓁* = 0.25, corresponding to a mean lag time of 4 days before ctDNA release from doomed irradiated cells. **(b):** Example linear regression fitting to patient data in nasopharyngeal carcinoma from Lo et al. [14] to the mean model-predicted ctDNA level (Appendix 6.4). **(c), (d):** Example simulations for varied survival rates and lag times. In (c), the survival rate per fractionated dose *S* is varied from 0.6 to 0.9. In (d), the lag time is varied from 3 to 7 days. The initial parameters are the same as in(a) otherwise.

The kinetics of ctDNA under radiation treatment are summarized in a review by McLaren and Aitman [17] and generally agree with predictions from our mechanistic model. As in targeted therapy and chemotherapy, radiotherapy patients have exhibited transient rises in ctDNA followed by eventual decreases within two weeks of treatment initiation [14, 29, 19]. However, in contrast to the ctDNA peaks observed at 26 hours in targeted therapy [27], peak ctDNA levels from radiation treatment has generally been observed 3-6 days after the first treatment in both xenograft models [29] and human nasopharangyeal carcinoma patients [14].

## 3 Biomarker design using virtual patient cohorts

We next use our models of treatment response to propose and evaluate various candidate biomarkers of response to each therapy. We focus on the goal of rapid prediction of therapy response using early time point ctDNA analyses. Table 1 reviews existing ctDNA biomarker analyses, indicating the potential of ctDNA assays for prognostic predictuion but also differences in biomarker design and efficacy across cancer and treatment types. Current ctDNA biomarker analysis is limited by a lack of higher time resolution data within the first few weeks of treatment and the focus on decrease in ctDNA alone as a prognostic indicator. A few case studies with daily ctDNA sampling during the beginning of treatment exhibit interesting features such as transient ctDNA peaks in responsive patients [27, 33, 14]. More sophisticated biomarker analysis has the potential to leverage these features for making personalized and rapid treatment decisions. However, no such data is currently available for larger groups of patients. Thus, here we utilize the models developed in Section 2 to simulate randomized virtual patient cohorts with higher time resolution early ctDNA sampling, for the development of treatment mechanism-specific biomarkers that predict treatment response.

### Targeted therapy biomarkers

The targeted therapy model from Section 2.1 was used to generate a randomized cohort of 1500 patients treated with targeted cytotoxic treatment. Details of the randomized cohort generation are provided in Section 6.1. We simulated the a dense data collection protocol, taking three evenly-spaced ctDNA samples in the 24 hours immediately before and three samples immediately after the start of treatment, and explored the development of biomarkers utilizing these sampled data to predict treatment response.

In particular, we consider two primary components as building blocks for biomarker design: *f*_*pre*_, the slope of the ctDNA signal immediately prior to the start of treatment, and *f*_*post*_, the slope of the ctDNA signal immediately after the start of treatment (see Figure 4a). These components were then combined in several different candidate biomarkers (Figure 4b) and assessed for predictive ability. These candidates were assessed based on their correlation strength with clinically relevant metrics. In particular, we considered the proportion of sensitive cells (PSC) at the start of treatment to be clinically predictive of long-term treatment failure, and the maximum tumor shrinkage (MTS) percentage to be an indicator of the depth of response during the treatment course. Scatterplots in figures 4c and 4d show the correlations between each biomarker candidate and MTS or PSC, and the table in Figure 4b summarizes the correlation coefficients. The most effective biomarker in this analysis was *V*_1_, which is a metric for the height of the jump in ctDNA levels upon initiation of treatment. These results suggest that larger height of the ctDNA peak induced by the start of targeted cytotoxic therapy should be correlated with both a stronger initial tumor burden reduction as well as long term treatment success.

**Figure 4:**
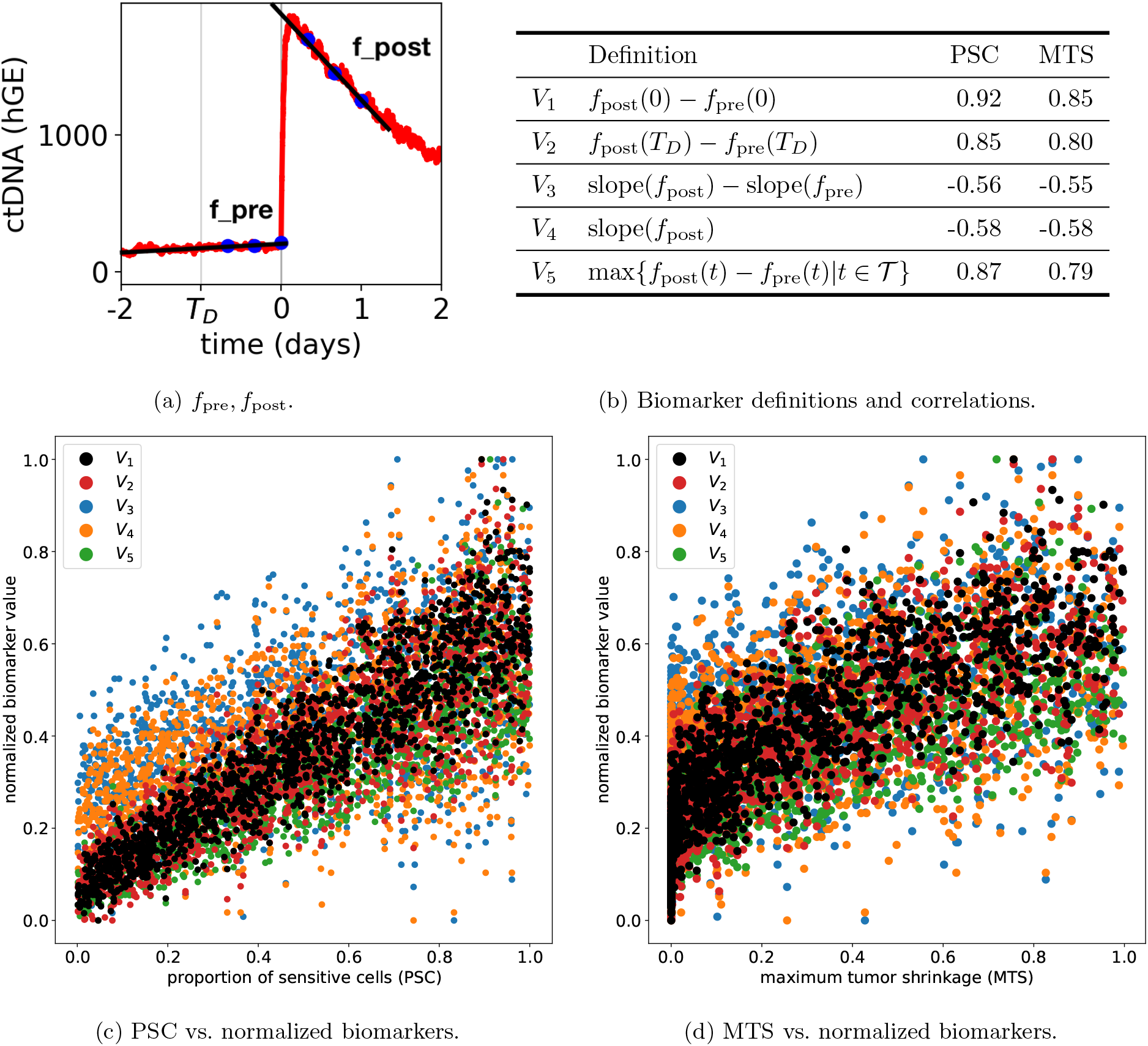
Biomarker definitions and analysis for targeted therapy. **(a):** Example diagram for *f*_pre_, *f*_post_. We fit lines *f*_pre_, *f*_post_ to the samples before and after initiation of treatment, respectively (blue dots). **(b):** Table of biomarker definitions and correlation coefficients of biomarkers and the initial proportion of sensitive cells (PSC) and the maximum tumor shrinkage (MTS), i.e. the largest proportional decrease in tumor burden, for a cohort of 1500 simulated patients with random parameters. The time of detection is denoted *T*_*d*_ and is equal to −1 in this simulation. The set of sample times after the start of treatment are denoted *T* . **(c), (d):** Correlations of PSC and MTS with normalized biomarker values for simulated cohort. Each biomarker was scaled to have values within [0, 1].

### Radiation therapy markers

The radiation therapy model from Section 2.2 was used to generate a randomized cohort of 1000 patients treated with radiation. Details of the randomized cohort generation are provided in Section 6.1. The patients were treated with a four week schedule of daily weekday radiation doses starting on day 7. ctDNA samples were collected on Monday, Wednesday, and Friday of the first two weeks of treatment (days 7, 9, 11, 14, 16, and 18). This extended sampling window, in comparison to targeted therapy, is motivated by effects of the lag time between radiation and apoptosis of lethally-irradiated cells, observed clinically [14] and in simulations (Figure 3d).

Using this simulated cohort data, several candidate biomarkers were explored using features of the sam-pled ctDNA dynamics – including the minimum and maximum sample levels, regression line slope, and weekly integrated ctDNA (the sum of ctDNA levels for each week). Figure 5 provides definitions of candidate biomarkers, which were evaluated by examining their correlation with the survival probability *S* of each cell after each fractionated dose of radiation (which measures efficacy of treatment), and the maximum tumor shrinkage (MTS) in the simulated cohort. By this measure, the most predictive biomarker was *R*_1_, which is an approximation of the area under the curve in the second treatment week divided by the area under the curve of the first week.

**Figure 5:**
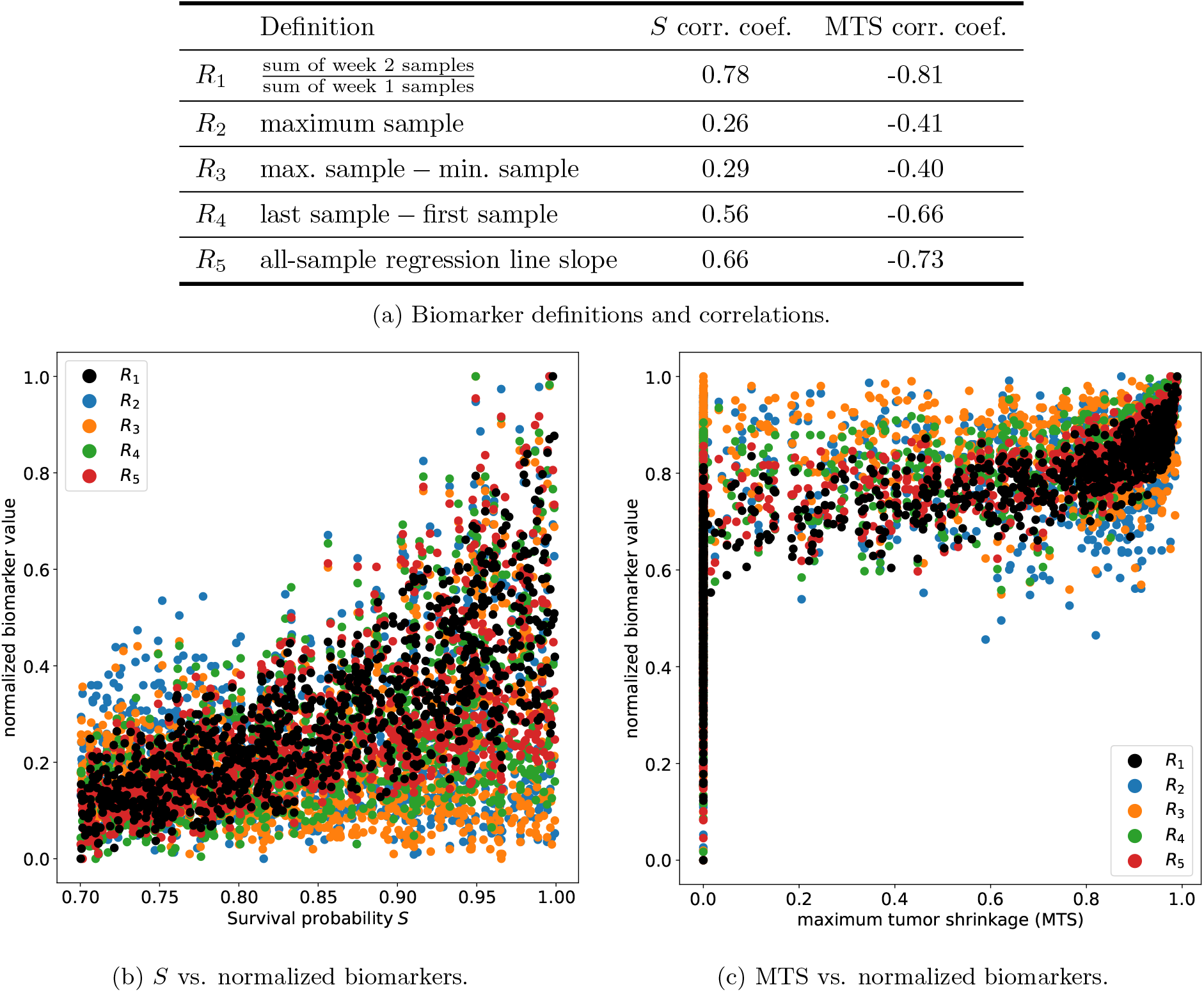
Biomarker definitions and analysis for radiotherapy. **(a):** Table of biomarker definitions and correlation coefficients of biomarkers with *S*, the survival probability for each fractionated dose and the maximum tumor shrinkage (MTS), i.e. the largest proportional decrease in tumor burden, for a cohort of 1000 simulated patients with random parameters. ctDNA samples were collected on Monday, Wednesday, and Friday of the first two weeks of treatment (days 7, 9, 11, 14, 16, 18). **(b), (c):** Correlations of S and MTS with normalized biomarker values for simulated cohort. Each biomarker was scaled to have values within [0, 1].

### Biomarker performance in predictions of partial response

To further evaluate the performance of the top candidate biomarkers (*V*_1_, *R*_1_), we assessed their ability to predict whether simulated patients would achieve partial response (PR), i.e., at least a 20% decrease in tumor burden during treatment (MTS *≥* 0.2). Figure 6 shows the receiver operating characteristic (ROC) curves for detection of partial response in the simulated cohorts for patients undergoing targeted cytotoxic therapy, chemotherapy and radiation. In each case, the top candidate biomarkers showed strong ability to predict PR. Using *V*_1_ with the simulated targeted therapy cohort data resulted in a ROC curve (Figure 6a) with an area under the curve of 0.92 for detection of partial response. In particular, the optimal threshold of *V*_1_ > 76 predicted PR with 91% sensitivity, 88% specificity, and 87% positive predictive value (PPV). The *V*_1_ biomarker was also evaluated as predictor of PR for a randomly simulated cohort of 1000 chemotherapy patients. We found that this produced an ROC curve with AUC 0.84 (Figure 6c). At the optimal threshold of *V*_1_ > 26, PR was predicted with 80% sensitivity, 74% specificity, and 61% PPV. Lastly, *R*_1_ was evaluated as a classifier of partial response in a simulated radiotherapy cohort of 1000 patients; this produced an ROC curve with AUC 0.96 (Figure 6b). The optimal threshold was found to be *R*_1_ ≤ 1.15, which predicted PR with 89% sensitivity, 93% specificity, and 95% PPV. Overall, the top-performing biomarkers (*V*_1_, *R*_1_) for targeted therapy, chemotherapy and radiotherapy demonstrated strong performance in predicting partial therapeutic response.

**Figure 6:**
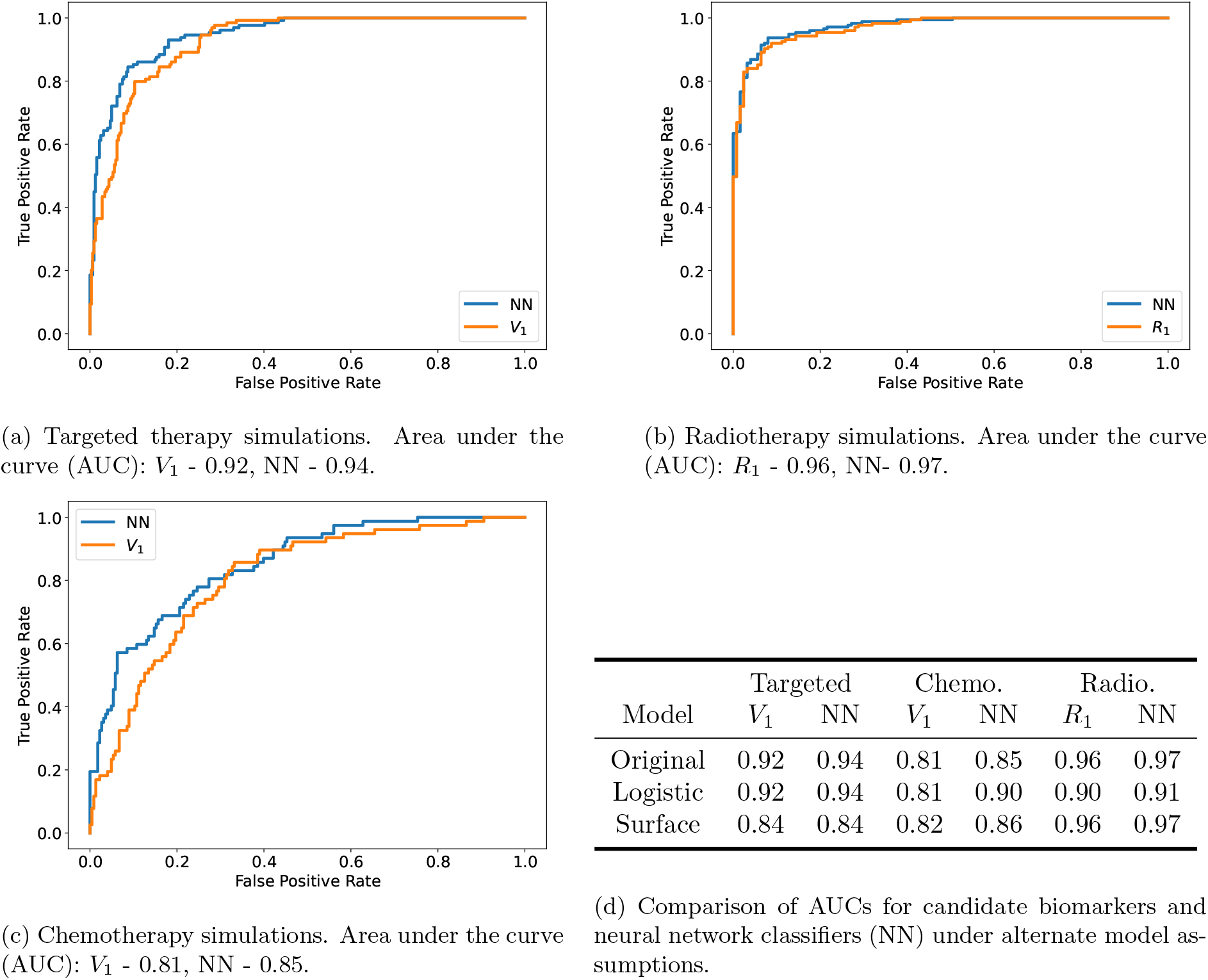
Detection of partial response (tumor shrinkage of at least 20%) in simulated random cohorts. **(a)-(c):** receiver operating characteristic (ROC) curves for prediction of partial response from simulated ctDNA data for candidate biomarkers (*V*_1_, *R*_1_) and for neural network classifiers (NN). ROC curves are shown for (a) 1500 simulated targeted therapy patients, (b) 1000 simulated radiotherapy patients, and (c) 1000 simulated chemotherapy patients. **(d):** In addition to the simulations under the original model assumptions, simulated cohorts for each treatment type were generated with models accounting for logistic growth with a carrying capacity and the assumption of spherical tumors with only surface cells shedding ctDNA.

### Comparison with neural network classifiers and model generalizations

A natural question that may arise is whether the metrics we have defined are optimal, i.e. are there features not captured by our biomarkers that would improve prediction of treatment response? In order to address this question, we trained and tested a PyTorch neural network classifier on the same randomized cohorts used for the biomarker analyses. The classifier was designed to receive the same inputs as the metrics and produce a binary prediction of whether the patient would exhibit partial response (PR) or not. Remarkably, we found that the performance of our biomarkers was able to match that of the neural network classifier. Figure 6 shows the ROC curves for our biomarkers for targeted therapy, chemotherapy, and radiotherapy in comparison with those of the neural network classifier. In all three treatments, the AUC for our metric matches that of the classifier extremely closely. This suggests that the metrics we have defined are optimal or near optimal and would not be significantly outperformed by alternate definitions. Moreover, it demonstrates that we do not have to sacrifice interpretability for performance.

Additionally, we assessed the performance of our biomarkers against the neural network classifier under alternate model assumptions (Appendix 6.2). In our model, we assume that the tumor is small enough relative to carrying capacity that its effects on growth are not relevant and that ctDNA shedding can originate from any cell. In order to analyze the validity and robustness of our observations, we tested our metrics using simulations under logistic growth with a carrying capacity and surface-only shedding. In both cases, *V*_1_ was still the best metric for targeted therapy and chemotherapy and *R*_1_ was still the best metric for radiotherapy. Moreover, in all scenarios, our biomarker analysis performed similarly to the neural network classifier (Figure 6d). This suggests that our observations are not overly sensitive to our model assumptions and in fact have predictive power under more general assumptions.

## 4 Discussion

This study leverages mechanistic modeling of therapeutic response and simulated randomized patient cohorts to develop ctDNA longitudinal biomarkers for predicting prognosis in tumors treated with targeted therapy, chemotherapy and radiation therapy. In the case of cytotoxic targeted therapy, we found that *V*_1_, a metric quantifying the height of initial ctDNA peaks, successfully predicted future partial response with 91% sensitivity, 88% specificity, and 87% PPV at the optimal threshold of *V*_1_ *≥* 76. For radiotherapy, the metric *R*_1_, an approximation of the area under the curve in the second treatment week divided by the area under the curve of the first week, predicted partial response with 89% sensitivity, 93% specificity, and 95% PPV at the optimal threshold of *R*_1_ *<* 1.15. For chemotherapy, the optimal threshold was *V*_1_ > 26, which predicted future partial response with 80% sensitivity, 74% specificity, but only 61% PPV, which may be driven by a lower number of cell deaths in comparison to successful targeted cytotoxic treatment. In all three treatments, we were able to define and validate dynamic early ctDNA biomarkers that performed favorably compared to existing biomarkers (Table 1) in terms of accuracy and earliness. These results suggest that early ctDNA kinetics with sufficient time resolution have the potential to provide valuable predictions of clinical outcomes.

Our proposed biomarkers rely upon frequent ctDNA samples collected within a short time window surrounding the initiation of therapy. In particular, for targeted therapy or chemotherapy, the biomarker *V*_1_ utilizes six samples collected during a 48-hour time period (24-hours prior to 24-hours after initiation of therapy. For radiotherapy, the quantity *R*_1_ utilizes six samples collected during the first two weeks after initiation of radiotherapy. These proposed sampling approaches differ significantly from most current sampling protocols in clinical practice, where ctDNA is collected at time points occurring several weeks or months apart, and sometimes does not commence until four weeks post therapy induction [27, 37, 5, 8]. Our results suggest that early-treatment ctDNA signals may hold valuable information about biological response to therapy. The rapid time scales of ctDNA production and decay, which are driven by biological and chemical processes occurring on the scales of hours and minutes, provide additional motivation for dense, frequent sampling within this early treatment period. Valuable prognostic information about tumor response to therapies could be overlooked under sampling protocols that are insufficiently dense or that miss the therapy induction window.

The current study establishes a general framework for the design of treatment-specific biomarkers for guiding the sampling and interpretation of ctDNA analyses, based on mechanistic modeling and the generation of virtual randomized patient cohorts. However, comparison with clinical observations, once available, is critical for evaluating and refining the specific biomarkers and sampling protocols proposed in this study. Limitations of the current work also suggest several directions for further study. For example, our current models of treatment response utilize a simple model of exponential tumor population growth. While we have demonstrated in this work that our biomarker performance is robust to a a few alternate growth models such as surface-driven and logistic growth, further exploration of more complex growth scenarios, spatial effects and pharmacokinetics is warranted. In addition, incorporating multidimensional ctDNA data tracking the clonal frequencies of various tumor-associated mutations is the subject of current work, as is the consideration of ctDNA dynamics under further treatment types and combination therapies.

## 5 Acknowledgements

J.F. was partially supported by NSF DMS 2052465, NSF CMMI 2228034, NIH R01CA241134, and Research Council of Norway Grant 309273. A.L. was partially supported by NSF DMS 2052465. K.L. was partially supported by NSF CMMI 2228034.

## 6 Appendix

### 6.1 Simulations of patient cohorts

We generated a cohort of 1500 random targeted therapy patients. As a simplifying assumption, we assume that the sensitive and resistant cells have the same birth and death rates off treatment. Following parameter estimates in [20], for each patient we uniformly generate a random birth rate *b*_*s*,1_ = *b*_*r*,1_ ∈ (0.1, 0.2) and death rate *d*_*s*,1_ = *d*_*r*,1_ ∈ (0.05, 0.0.08). We also randomly generate an initial population *N ∈* (8000, 12000), initial proportion of sensitive cells in (0, 1), and detection size in (*N* + 2500, *N* + 5000). When the total tumor population reaches the detection size, we collect three samples in the 24 hours immediately before the initiation of treatment and three samples in the 24 hours immediately after initiation of treatment. When treatment is started, the birth and death rate of the sensitive cells become *b*_*s*,2_ ∈ (*b*_*s*,1_ − 0.05, *b*_*s*,1_) and *d*_*s*,2_ ∈ (*d*_*s*,1_ + 0.3, *d*_*s*,1_ + 0.5). Here we make another simplifying assumption that the resistant cells are fully resistant and do not change their birth and death rate on treatment.

To simulate a chemotherapy cohort, we simulated 1000 patients with the same parameter ranges. However, on treatment we defined an efficacy rate *K*_*s*_ in (0.7, 1.0) and *K*_*r*_ = 0, again assuming that the resistant cells are fully resistant.

For our radiotherapy cohort, we started with the same underlying patient tumor characteristics. The simulated patients followed a four week treatment schedule of daily radiation doses from Monday to Friday. For each patient, we generated a survival rate *S* in (0.7, 1.0) that represented the proportion of tumor cells that survived each dose. We also generate a lag parameter *𝓁* in (0.2, 0.3) to capture the 3 − 5 day lag time described by [17].

### 6.2 Model validation

To create our neural network classifier, we defined a model with three layers each with 60 nodes. We trained the model for 300 epochs on 70% of the data and reserved the rest of the data for testing. The model received the six ctDNA samples as well as a binary classification of whether partial response was achieved or not.

To test variations in the model assumptions, we simulated 1000 patients for each targeted therapy, chemotherapy, and radiotherapy using the same parameter ranges as the original experiments, but using logistic growth with a carrying capacity *K ∈* (20000, 40000) uniform for the tumor growth. We also simulated cohorts assuming that only cells on the surface of the tumor produced ctDNA. We assumed a spherical tumor and that there were 10^8^ cells per cubic centimeter [18, 21].

### 6.3 Summary statistics of targeted therapy and chemotherapy models

**Proposition 1**. *Let C*(*t*) *denote the ctDNA quantity in the bloodstream at time t with treatment starting at time 0. Assuming that C*(0) *is a Poisson random variable with mean* 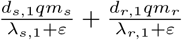 *we have that for targeted therapy and chemotherapy*,

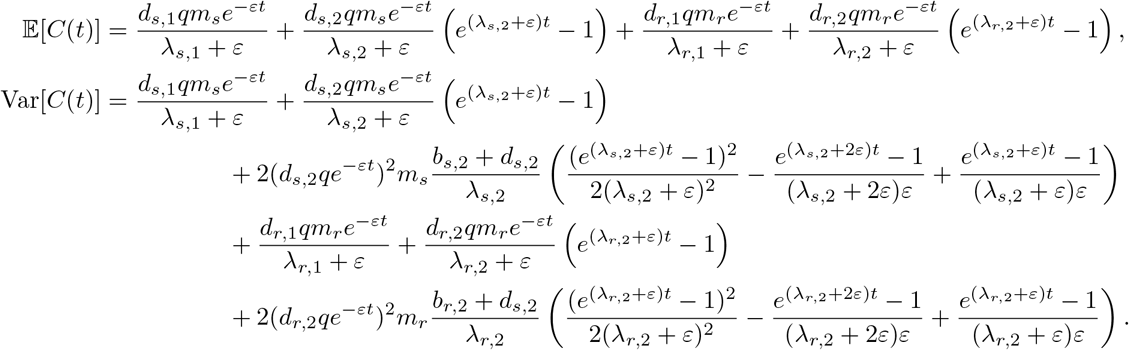

*Proof*. Consider the contribution from pre-treatment ctDNA from the sensitive cells, *C*_*s*,pre_(*t*). Conditioning on the sensitive population size at time 0 being *m*_*s*_, we have that

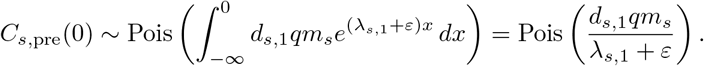

Applying the thinning property with the shedding probability *ε*, we have that

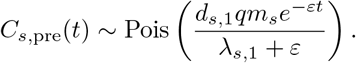

Now consider *C*_s, post_(*t*). We know that given {*N*_*s*_(*x*)}_*x≤t*_,

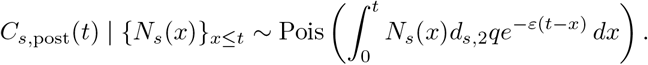

Then

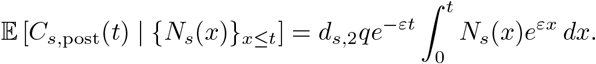

Thus,

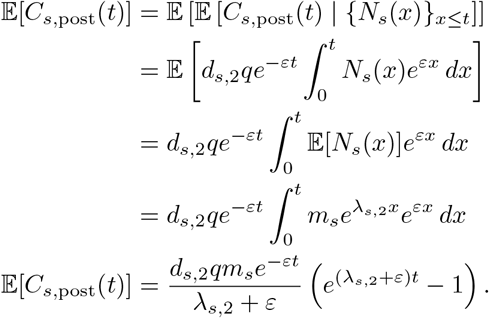

Then we have that

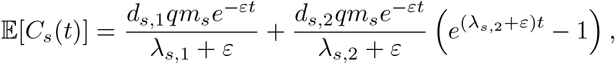

Similarly, the expected value for the resistant population is

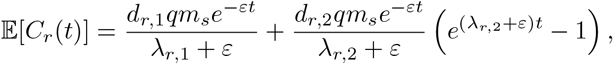

so we are done.

Now let us consider the variance. By the law of total variance, we have that

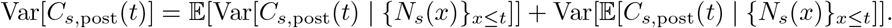

Since *C*_*s*,post_(*t*) | {*N*_*s*_(*x*)}_*x*≤*t*_ is Poisson distributed, we have that

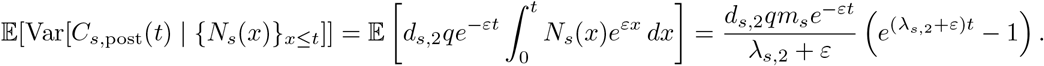

Now consider

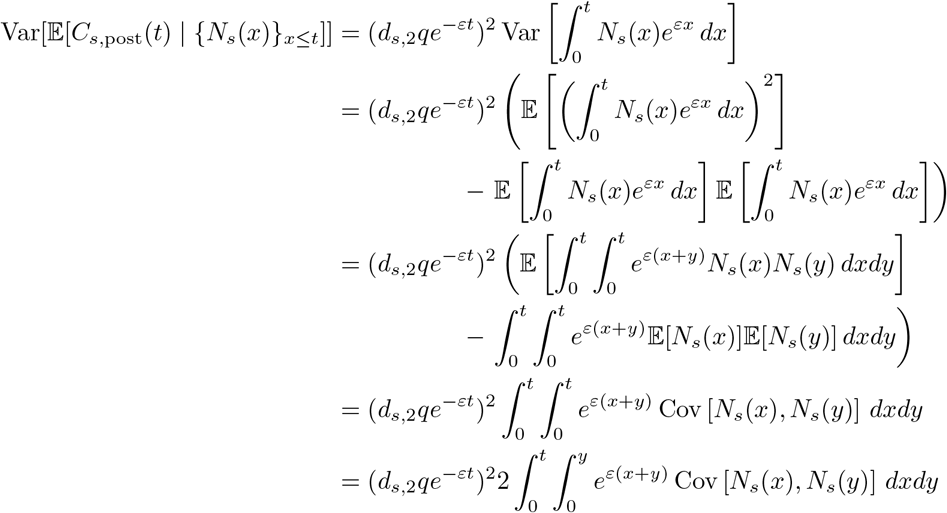

Assume 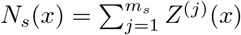 f or i.i.d. stochastic processes {*Z*^(*j*)^(*x*) : *x ≥* 0, *j ≥* 1}. Thus,

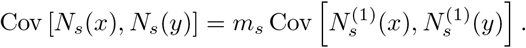

Notice that

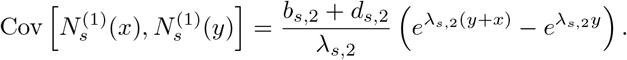

So

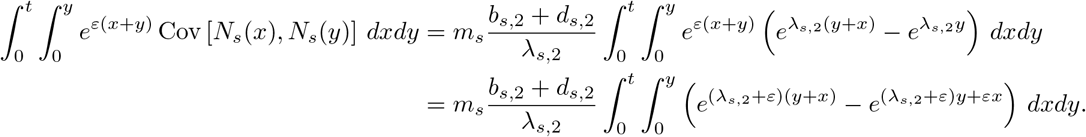

Since

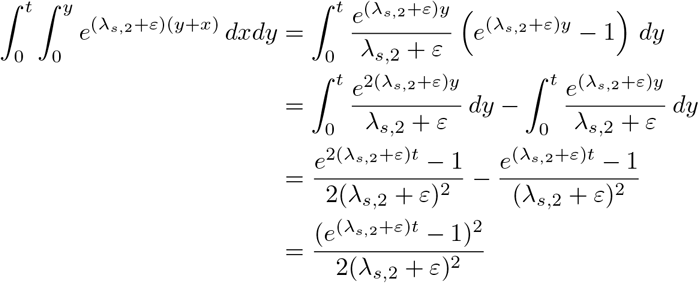

and

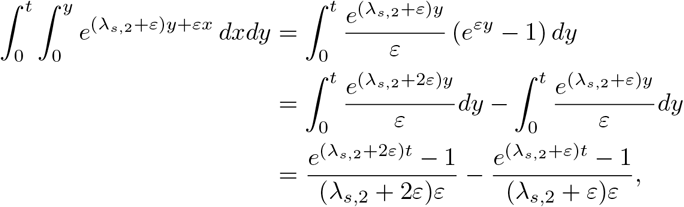

we have that

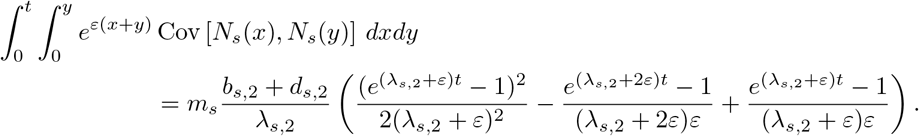

Thus,

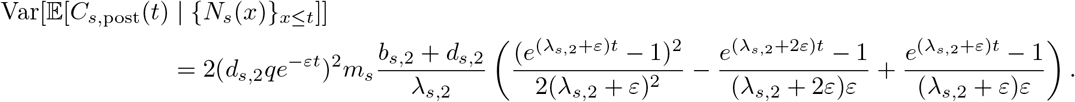

So we have that

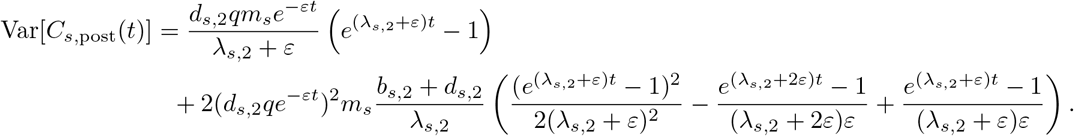

Then

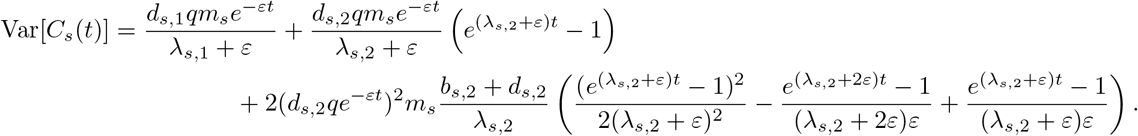

The same applies to Var[*C*_*r*_(*t*)] so we are done.

### 6.4 Summary statistics of radiotherapy model

#### Lemma 1.

*Let N* (*t*) *be the healthy tumor population at time t and let radiotherapy treatments be applied under our model at times {t*_*1*_ , *t*_*2*_ , … , *t*_*n*_ *} with dosage D Gy and survival probability* 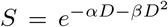. *Let k* = max{*i*|*t*_*i*_ *< t}. Then*

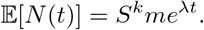

*Proof*. Proceeding by induction, we will show that

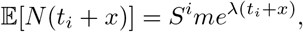

for all *x* such that *t*_*i*_ *< t*_*i*_ + *x ≤ t*_*i*+1_. Assume that the previous statement holds for some fixed *i* a nd now consider 𝔼[*N* (*t*_*i*+1_ + *y*)] for *t*_*i*+1_ *< t*_*i*+1_ + *y ≤ t*_*i*+2_. We know that if *t*_*i*_ + *x* = *t*_*i*+1_,

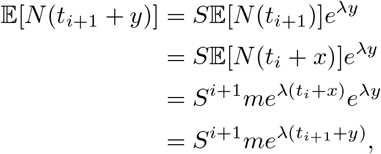

as desired. As a base case we know that for *t ≤ t*_1_ 𝔼[*N* (*t*)] = *me*^*λt*^. So by induction, we have that for all *t >* 0, if *t* = *t*_*k*_ + *x* where *t*_*k*_ = max{*t*_*i*_|*t*_*i*_ *< t}*, then

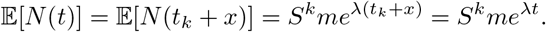

#### Proposition 2.

*Let C*(*t*) *be the amount of ctDNA present at time t in the above scenario. Then*

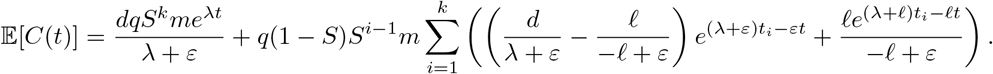

*Proof*. By Lemma 1, if *k* = max{*i*|*t*_*i*_ *< t}* then 𝔼[*N* (*t*)] = *S*^*k*^*me*^*λt*^. Consider the cells killed by the *i*-th radiation treatment, *I*_*i*_(*t*). We know that for *t ≥ t*_*i*_,

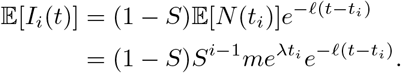

Now let us consider *C*(*t*). Denote the ctDNA contribution of lineages that are killed by the *i*-th treatment as *C*_*i*_(*t*). ctDNA from cell lineages that survive until time *t* will be denoted *C*_*∞*_(*t*).

Then by Lemma 1,

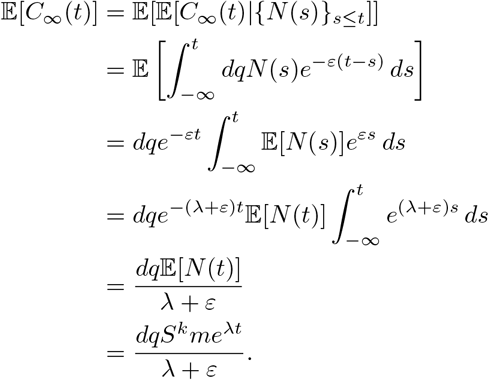

Now consider *C*_*i*_(*t*), the ctDNA contribution of lineages killed by the *i*-th treatment. Let *C*_*i*,pre_(*t*) be the ctDNA shed by the lineages before the radiotherapy treatment and let *C*_*i*,rad_(*t*) be the ctDNA shed from the decay of irradiated cells. Then

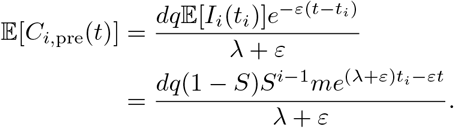

And

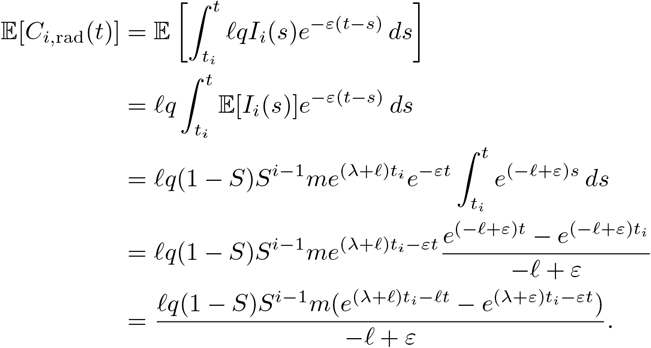

Then all together we have that

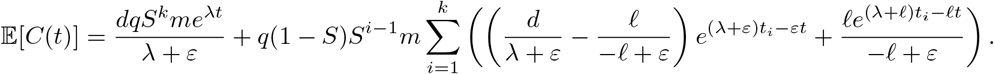

